# What’s in the box? Authentication of *Echinacea* herbal products using DNA metabarcoding and HPTLC

**DOI:** 10.1101/202721

**Authors:** Ancuta Cristina Raclariu, Carmen Elena Ţebrencu, Mihael Cristi Ichim, Oana Teodora Ciupercǎ, Anne Krag Brysting, Hugo de Boer

## Abstract

**Background:** Differences in regulatory policies between countries as well as a lack of appropriate standardized methods for authentication and quality control of herbal products lead to concerns over quality and safety. *Echinacea* products are among the top selling herbal products in Europe and the United States with indications for broad range of ailments.

**Purpose:** This study approached the need for a novel analytical strategy in authentication of herbal products.

**Methods:** A combination of high performance thin layer chromatography (HPTLC) and DNA metabarcoding was employed. Fifty-three *Echinacea* herbal products marketed across Europe were tested to evaluate the accuracy of these methods in plant identification and their potential in detecting substitutes, adulterants and other unreported plant constituents.

**Results:** HPTLC provides a high resolution in detecting *Echinacea* phytochemical target compounds, but does not offer information on the other species within the product. Alternatively, we showed that the limitation of HPTLC to detect non-targeted species can be overcome through the complementary use of DNA metabarcoding. Using DNA metabarcoding, *Echinacea* species were detected in 34 out of the 38 retained products (89 %), but with a lack of discriminatory resolution at the species level due to the low level of molecular divergence within the *Echinacea* genus. All herbal products showed considerable discrepancies between ingredients listed on the label and the ones detected using DNA metabarcoding registering an overall ingredient fidelity of 43 %.

**Conclusion:** The results confirm that DNA metabarcoding can be used to test for the presence of *Echinacea* and simultaneously to detect other species present in even highly processed and multi-ingredient herbal products.

**Abbreviations:**

EMA: European Medicines Agency
BLAST: basic local alignment search tool
FTIR: Fourier-transformed infrared spectroscopy
IR: infrared spectrometry
HPTLC: high performance thin layer chromatography
*mat*K: maturase K
MS: mass spectrometry
MOTU: molecular taxonomic unit
nrITS: nuclear ribosomal internal transcribed spacer
Ph.Eur.: European Pharmacopoeia
*rbc*L: ribulose bisphosphate carboxylase
TLC: thin-layer chromatography
UV-VIS: ultraviolet-visible

## 1. Introduction

### 1.1 Legislation and *Echinacea*

The regulation of herbal products varies globally, and products can be classified as food supplements, medicines, homeopathic products, cosmetics or even biocides depending on specific legislation. The European Medicines Agency (EMA) regularly produces updates of the European Pharmacopoeias (EDQM, 2014) and in addition has produced a number of monographs on quality and authentication of specific herbals providing relevant methodological specifications for their quality assessment. However, the primary legal responsibility for the safety of the marketed products is delegated by default to the manufacturers.

Sales of herbal dietary supplements have seen year-on-year increases of 5-10 %, and in 2014 sales reached an estimated total of more than $6.4 billion, with *Echinacea* among the top five top grossing taxa (Smith et al., 2015). *Echinacea* is a genus of composites (Asteraceae) comprising a small number of herbaceous grasslands perennial taxa whose natural distribution is limited to North America (Kindscher and Wittenberg, 2016). Three *Echinacea* species, *E. angustifolia* DC. (Narrow-leafed purple coneflower), *E. pallida* (Nutt.) Nutt. (Pale purple coneflower) and *E. purpurea* (L.) Moench (Purple coneflower), are used in traditional herbal medicine (EDQM, 2014).

Today *Echinacea* is cultivated widely in Europe and North America for use in commercial herbal products in a diverse range of products, including herbal teas, capsules, tablets, powders, tinctures and beverages (Brown et al., 2011). A 2007 survey by the National Centre for Complementary and Alternative Medicine showed that *Echinacea* was the most commonly used herbal medicine among adults and children in United States (Barnes et al., 2008). In Europe, a study on the use of food supplements and medicines showed that *Echinacea* and *Ginkgo* are the most common herbals (EAS, 2006).

### 1.2 Pharmacological use and effects of *Echinacea* species

*Echinacea* has been extensively studied for its pharmacological effects in both *in vitro* and *in vivo* studies (Table S1). *In vivo* human clinical studies on the oral administration of *E. purpurea* commercial herbal juice have failed to confirm previous *in vitro* findings (Schwarz et al., 2005, 2002; Sperber et al., 2004). However, a recent study conducted on 68,522 Norwegian women and their children revealed no risk of malformations or adverse pregnancy outcomes associated with the use of *Echinacea* in pregnancy (Heitmann et al., 2016).

### 1.3 Chemical profiles and differences between *Echinacea* species

The constituents of *Echinacea* species used for medicinal purposes differ qualitatively and quantitatively (Barnes et al., 2005; Binns et al., 2002). Alkamides, polysaccharides, glycoproteins, volatile oils and phenolic compounds have been considered important constituents of the plant (Table S2). *Echinacea purpurea* shows a similar phenolic phytochemical profile (cichoric acid, caftaric acid, chlorogenic acid) in roots, flower and leaves, but with no cynarine in the aerial parts, whereas *E. angustifolia* roots shows low amounts of cichoric acid and cynarine, and that of *E. pallida* shows both cichoric acid and cynarine. *Echinacea purpurea* aerial parts contain no echinacoside, while echinacoside is a major component in the roots of *E. pallida* and *E. angustifolia* (and also present in small quantities in the flower and leaves Alkamides are found in the rhizomes and roots of *E. angustifolia*, and less abundantly in the aerial parts of *E. purpurea* and mainly absent in *E. pallida* roots). *E. pallida* contains large amounts of ketoalkenes. Rutoside is a flavonoid present in the leaves of all the three species of *Echinacea* (Barnes et al., 2005).

### 1.4 Quality issues of *Echinacea* herbal products

The increased use of *Echinacea* species has led to concerns about adulterated products resulting from challenges in morphology-based identification, due to overlapping morphological variability and frequent hybridization between species (Flagel et al., 2008). Furthermore, reported adulteration of *E. purpurea* with the roots of *Parthenium integrifolium* L., *Lespedeza capitata* Michx., *Eryngium aquaticum* L., *Rudbeckia nitida* Nutt., *Helianthus annuus* L. or *Liatris aspera* Michx. lead to safety concerns of the herbal products (Zhang et al., 2017). The use of unreported ingredients is a serious safety concern as adverse drug reactions cannot be associated to the product label and ingredients (Gilbert, 2011). Commercially available herbal products contain one or more *Echinacea* species originating from the same or different geographical areas, and the resulting phytochemical diversity across these products can complicate further investigations (Barnes et al., 2005). *Echinacea* secondary metabolites are used for the qualitative identification of species in the industry. Presence of for example, cichoric acid for *E. purpurea and E. angustifolia*, echinacoside for *E. angustifolia* and *E. pallida*, ketoalkene for *E. pallida* were routinely tested (Mistrikova and Vaverkova, 2006). Nevertheless, the presence of these markers does not provide unequivocal identification of the species since, for instance, traces of echinacoside can be also found in *E. purpurea* (Arnason et al., 2002). Presently, the differentiation of species is based on the relative abundances of metabolites by various phytochemical techniques, but marketed herbal products are often highly processed complex formulations with numerous ingredients, and these methods might not enable accurate identification of all plant ingredients, especially if target species are admixed with other species (De Boer et al., 2015).

### 1.5 DNA metabarcoding of herbal products and *Echinacea*

High phenotypic plasticity within *Echinacea* has complicated the taxonomy of the genus and led to misidentifications, confused taxonomies and misapplication of taxa (Kindscher and Wittenberg, 2016). Combined plastid *(trnS* and *trnG*) and nuclear (*Adh*, *CesA*, and *GPAT*) phylogenies found no resolved topologies, suggesting incomplete lineage sorting, as well as the potential for widespread hybridization and backcrossing following secondary contact within the genus (Flagel et al., 2008). Flagel et al. (2008) note that in contrast to the low discriminatory resolution of these molecular markers, previous studies using morphological characters and metabolic profiles found well-resolved and taxonomically supported relationships within *Echinacea*, and suggest the use of markers with more broad genomic coverage. Zhang et al. (2017) recovered the complete plastid genomes from all nine *Echinacea* species, and based on both coding and non-coding regions of the chloroplast genomes, *Echinacea* species were separated into two clades with strong support. In addition, Zhang et al. (2017) report that the core barcoding markers *mat*K and *rbc*L do not differ sufficiently for species-specific identification of *Echinacea* but suggest a combination of nrITS and *trnH-psbA* as the optimal barcoding markers instead. The incongruence in nuclear ribosomal and plastid phylogenies, reported by Flagel et al. (2008), however suggests that neither plastid markers nor nrITS provide an accurate picture of the phylogenetic history of the genus.

In this study, we use DNA metabarcoding to detect species diversity in *Echinacea* herbal products. The use of DNA metabarcoding for the identification of commercialized plant products has evolved with advances in molecular biology and sequencing (Coghlan et al., 2012; Raclariu et al., 2017b), and is defined as high-throughput multispecies (or higher-level taxon) identification using the total and typically degraded DNA extracted from an environmental sample (Taberlet et al., 2012). Here we test the hypothesis that *Echinacea* species are frequently admixed with other species in *Echinacea* herbal products. We test this hypothesis by authenticating European *Echinacea* herbal products using DNA metabarcoding and HPTLC to authenticate these using phytochemical constituents, and aim to answer the following research questions: 1) Can DNA metabarcoding be used to test for the presence of *Echinacea* species in herbal products, and to detect the presence of off label plant species due to substitution or adulteration?; 2) Can HPTLC be used to distinguish *Echinacea* species and to identify its exclusive presence in herbal products?

## 2. Materials and Methods

### 2.1 Sample collection

Fifty-three herbal products listing different *Echinacea* species, marketed as single (35) and multi-ingredient (18) food supplements consisting in herbal teas (9 loose teas and 8 bagged teas), capsules (16), tablets (13) and extracts (7), were purchased from different retail stores (20), pharmaceutical companies (32) and via e-commerce (1). The countries of origin of these products were Austria (1), China (1), Czech Republic (3), France (1), Germany (4), Italy (2), Macedonia (1), Poland (2), Romania (26), Switzerland (1), United States (7), and four of the products did not specify a country of origin. A list of samples is included as Table S3, but the producer/importer name, lot number, expiration date and any other information that could identify the specific products and producers are omitted. These herbal products were imported into Norway for scientific analysis under Norwegian Medicines Agency license no. 16/04551-2.

### 2.2 High performance thin layer chromatography (HPTLC) analysis

*Echinacea* phenylpropanoids (echinacoside, cynarin, cichoric acid, chlorogenic acid, caffeic acid, caftaric acid) from 53 herbal products were analyzed using HPTLC following recommendations in the specific monographs from the European Pharmacopoeia 8^th^ Edition (EDQM, 2014) and the CAMAG application notes on HPTLC identification of *Echinacea* species (2015). Herbal products were processed depending on their pharmaceutical formulation. Capsules (about 300 mg), tablets (400 mg), and teas (500 mg) were homogenized and mixed in a 1:100 ratio with methanol/water (80:20 V/V) in an ultrasonic bath for 15 min, and tinctures were diluted in 1:3 methanol. The obtained solutions were centrifuged and the filtered supernatant was collected and used as sample for further analysis. The following chemical and botanical standards were used. *Reference substances:* caftaric acid (HPLC grade, Carl Roth GmbH + Co. KG, Karlsruhe, Germany), cynarine (min. 98.0 %, HPLC/Sigma-Aldrich Co, St. Louis, Missouri, USA), cichoric acid (min. 95.0 %, Sigma-Aldrich, St. Louis, Missouri, USA), echinacoside (min. 98.0 % HPLC, Sigma-Aldrich Co, St. Louis, Missouri, USA), chlorogenic acid (min. 95.0 %, Sigma-Aldrich), caffeic acid (TLC grade, Carl Roth GmbH + Co. KG, Karlsruhe, Germany), verbascoside (min. 99.0 % HPLC, Extrasynthese SAS, Lyon, France), ferulic acid (min. 99.0 %, Sigma-Aldrich, St. Louis, Missouri, USA). *Solvents*: ethanol, min. 96 % V/V, and methanol, 99.3 % analytical reagent (Chimreactiv SRL, Bucharest, Romania). *Analytical reagents*: formic acid ACS (Merck KGaA, Darmstadt, Germany), ethyl acetate (min. 99.5 %, Sigma-Aldrich Co, St. Louis, Missouri, USA), distilled water prepared in laboratory; natural products - polyethylene glycol (NP/PEG) reagent (Honeywell Fluka, Seelze, Germany). *Botanical standards:* United States Pharmacopoeia (USP) analytical reference botanical standards: USP Powdered *Echinacea purpurea* F0D018, USP Powdered *Echinacea angustifolia* G0I377 and USP Powdered *Echinacea pallida* F0I285 (Rockville, Maryland, USA). *Apparatus*: A CAMAG high performance thin layer chromatography system (CAMAG AG, Muttenz, Switzerland), with a Linomat IV sample applicator, a Canon digital camera, and the following software, Reprostar III with winCATS planar chromatography manager software, Digistor II digital system with winCATS software with an Image Comparison Viewer enabling high resolution visualization of multiple samples for comparison of specific compound retention times and the included references.

Samples and references were analyzed on Silica gel HPTLC plates (60 F245 20×10 and 10×10 cm for tinctures and solutions, Merck KGaA, Darmstadt, Germany). The reference solutions were dissolved in methanol to obtain the following concentrations: caftaric acid 0.006 %, cynarine 0.02 %, cichoric acid 0.005 %, echinacoside 0.018 %, chlorogenic acid 0.015 %, caffeic acid 0.011 %, verbascoside 0.01 %, ferulic acid 0.01 %, USP Powdered *Echinacea purpurea* F0D018 1.0 %, USP Powdered *Echinacea angustifolia* G0I377 2.0 % and USP Powdered *Echinacea pallida* F0I285 1.0 %. *Echinacea* product sample extracts and reference solutions were applied separately in twin bands, on maximum 12 tracks at 4-8 mm distance using different application volumes (4-16 μl for extracts, 12-17 μl for bag and bulk teas, 12-17 μl for tablets, 9-15 μl for capsules and 3-6 μl for standard solutions) with a band length of 8-10 mm and a delivery speed of 8 s/μl using a CAMAG Linomat IV automatic sample applicator (see above). The plates were run for 30 min in the developing box and subsequently dried at 105 °C for 5 min. For phenylpropanoids identification, the plates were inspected at 254 and 366 nm before and after homogenous spraying with the Natural products-polyethylene glycol reagent (NP/PEG), and subsequently air dried. For each set of samples, the plates were developed in a saturated vertical-developing chamber at room temperature (20-22 °C) for 30 min with ethyl acetate:formic acid:acetic acid:water = 20:2.2:2.2:5.4 V/V as mobile phase; the development distance was 7 cm; after the development, the plates were air dried at room temperature. Derivatization of the chromatograms were performed by spraying the plates with NP (0.1 g in 10 ml methanol) and PEG 400 (0.5 g in 10 ml ethanol), followed by heating of the plates at 105 °C temperature for 15 min. Plates were subsequently imaged using a CAMAG Reprostar 3 with digital video camera at 254 nm for developed plates and 366 nm for derivatized plates. HPTLC densitometry (CAMAG- TLC Scanner 3 with WinCATS Planar Chromatography Manager Software) was done using spectra recording from 200-700 nm, absorption at 254 nm wavelength, slit dimension 8.00 x 0.40 mm, scanning speed 100 nm/s, and peak area evaluation by linear or polynomial regression.

### 2.3 DNA metabarcoding

The total DNA was extracted from small amounts (about 300 mg) of each homogenized herbal product or silica gel dried leaves, using the method described in Raclariu et al. (2017b). Amplicon DNA metabarcoding using barcoding markers nrITS1 and nrITS2 was done using the exact method described in Raclariu et al. (2017b), and sequenced on an Ion Torrent Personal Genome Machine (Life Technologies, Thermo-Fischer Scientific, USA). Sequencing read data was processed using the HTS barcode-checker pipeline (Lammers et al., 2014) as described in Raclariu et al. (2017b). A 99 % sequence similarity threshold was used for MOTU clustering with a minimum of 10 reads per cluster to reduce the formation of false MOTUs and the potential effects of sequencing bias known to affect the Ion Torrent sequencing platform (Loman et al., 2012). One representative sequence from each MOTU was taxonomically assigned using the Basic Local Alignment Search Tool (BLAST) (Altschul et al., 1990) against a reference nucleotide sequence database represented by a local copy of the NCBI/GenBank. BLAST results with a maximum e-value of 0.05, a minimum hit length of 100 bp, and similarity of > 99 % against the reference barcode were accepted as species level matches.

## 3. Results

### 3.1 High performance thin layer chromatography (HPTLC)

Identification and detection of *Echinacea* species, *E. purpure*, *E. angustifolia* and *E. pallida*, from tea, tablets, capsules, and extracts were done using HPTLC with the standard *Echinacea* phenylpropanoid references echinacoside, cichoric acid, caftaric acid, chlorogenic acid, and cynarine, and the botanical standards of *E. purpurea*, *E. angustifolia*, *E. pallida*. In the HPTLC chromatograms visualized at 366 nm after derivatization, the occurrence of fluorescent blue spots indicates the presence of phenylpropanes (Figure 1). The phenylpropanoid refraction (Rf) values were: 0.15-0.17 (echinacoside), 0.47-0.49 (chlorogenic acid), 0.51-0.53 (caftaric acid), 0.65 (cynarine), 0.87-0.9 (cichoric acid), 0.96-0.97 (caffeic acid), and other polyphenols (ferulic acid and verbascoside) at Rf = 0.35; 0.46; 0.52; 0.73 and 0.90.

**Figure 1.**
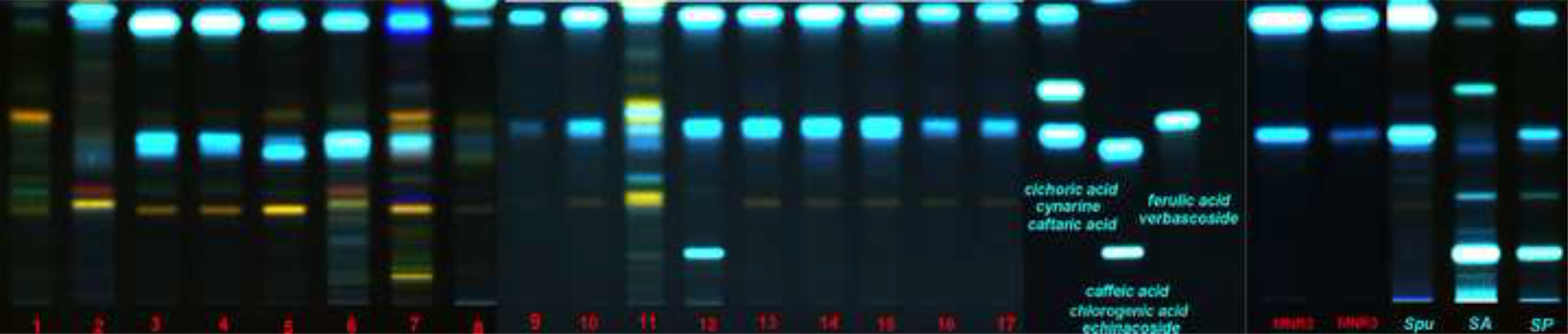
High performance thin layer chromatogram (HPTLC) based identification and detection of phenylpropanoids from the *Echinacea* sp. herbal products. The track assignment: (1) test solutions (1-53) (2) phenylpropanoids chemical reference solutions in the following order (increasing Rf): echinacoside, cholorogenic acid, caftaric acid, cynarin, cichoric acid, caffeic acid; (3) botanical reference solution of *Echinacea purpurea* (root (MNR1) and aerial parts (MH1)) (4) United States Pharmacopoeia (USP) analytical reference botanical standards (*E. purpurea* (Spu), *E. angustifolia* (SA) and *E. pallida* (SP)) **A**. HPTLC chromatograms of herbal teas. **B**. HPTLC chromatograms of capsules. **C**. HPTLC chromatograms of tablets. **D**. HPTLC chromatograms of extracts. The chromatograms were scanned after derivatization in UV 366 nm. Details about the herbal products can be found in the Table S3.

The results of HPTLC chromatographic identifications of the analysed herbal teas yielded the following results for the 10 samples labelled as including *Echinacea* sp. (Figure 1A and Table S4): Sample #1 doesn’t contain any of the targeted phenylpropanoids specific to *Echinacea* species; sample #8 has caffeic acid and low caftaric acid and content suggesting that only *E. purpurea* is present and in low concentration; sample #12 has very strong zones for echinacoside, cichoric acid and caftaric acid indicating a mixture of *E. purpurea* and *E. pallida*; seven samples (samples #4, #6, #9, #10, #11, #13, and #14) show distinct presence of caftaric and cichoric acid indicating presence of *E. purpurea*. The six samples labelled as including specifically *E. purpurea* yielded the following: in four samples (samples #3, #15, #16, #17) the presence of this species was confirmed by high content of caftaric and cichoric acid; in sample #2 low content cichoric and caffeic acid suggests a low concentration of *E. purpurea*; in sample #7 the presence of caftaric acid and a comparison of the profile with that of the botanical standard *E. angustifolia* (Figure 1A) indicates a mixture of *E. purpurea* and *E. angustifolia*. In sample #5 *E. purpurea* was identified from the distinct presence of cichoric and caftaric acid.

The results of the HPTLC chromatographic identifications of the analysed capsules yielded the following (Figure 1B and Table S4): The three samples labelled as including *Echinacea* sp. (#18, #31, #32) contained caftaric acid and cichoric acid, and this confirmed the presence of *E. purpurea*. In the nine samples labelled as including specifically *E. purpurea* (samples #19, #22, #23, #26, #27, #28, #29, #30, #33) the content of cichoric acid and caffeic acid confirmed the presence of this species, except for sample #29. In sample #21 labelled as including *E. angustifolia* the detection of echinacoside and cichoric acid and the comparison of the profile with that of the botanical standard *E. angustifolia* (Figure 1B) indicates the presence of a mixture of *E. angustifolia* and *E. purpurea*. In sample #24 labelled as including *E. pallida* the presence of echinacoside and cichoric acid and the comparison of the profile with that of the botanical standard *E. pallida* indicates the presence of a mixture of *E. pallida* and *E. purpurea* (Figure 1B). In samples #20 and #25 labelled as including mixtures of *E. purpurea* and *E. angustifolia*, the presence of these species was confirmed by the content of echinacoside, caftaric acid, cichoric acid and caffeic acid.

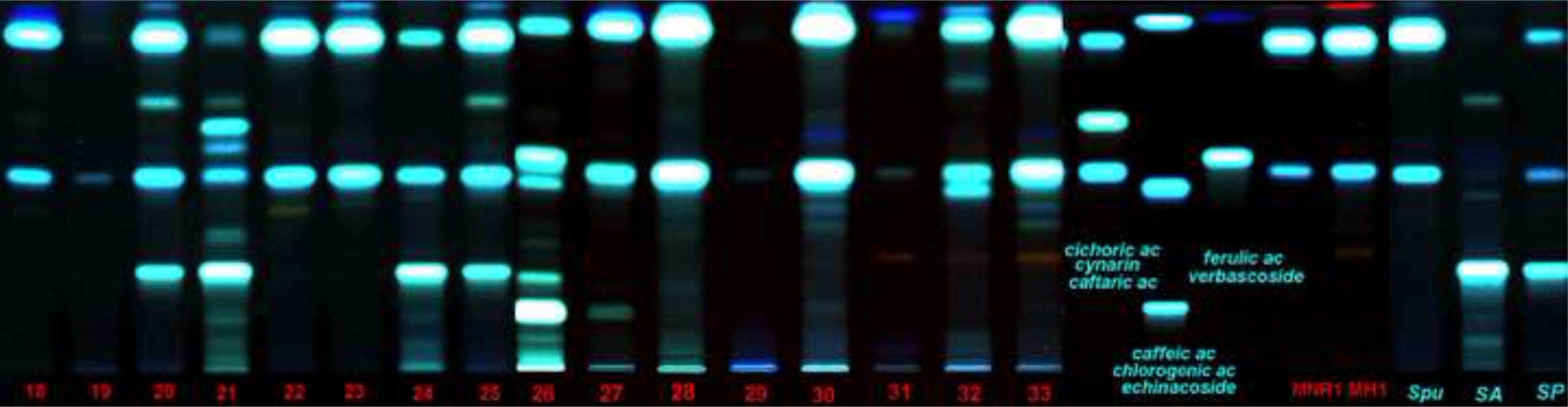

The results of the HPTLC chromatographic identifications of the analysed tablets yielded the following (Figure 1C and Table S4): The two samples labelled as including *Echinacea* sp. (samples #36, #46) differ in their chromatographic profiles. Sample #36 differs from that of the *Echinacea* species, and sample #46 contains caffeic acid and cichoric acid indicating the presence of *E. purpurea*. For the eleven samples labelled as including *E. purpurea* (samples #34, #35, #37, #38, #39, #40, #41, #42, #43, #44, #45) the presence of this species was confirmed in eight samples by the presence of caffeic acid and cichoric acid, and the presence of the echinacoside, caftaric acid and cichoric acid indicated a mixture of *E. purpurea* and *E. pallida* in sample #40, as well as a low concentration of *Echinacea* in sample #41. In sample #42 no *Echinacea* was identified.

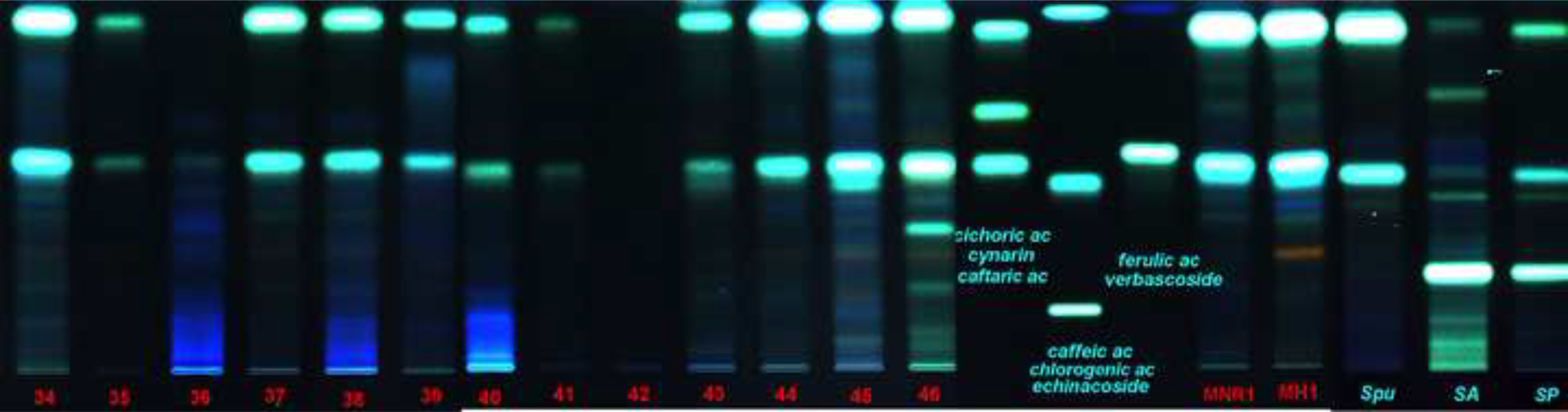

The results of the HPTLC chromatographic identifications of the analysed extracts yielded the following (Figure 1D and Table S4): Of the five samples labelled as including *E. purpurea* (samples #47, #49, #51, #52, #53), sample #47 was a glycerinated solution and this hampered an accurate composition estimation, sample #53 contained very weak zones for echinacoside, caftaric acid and cichoric acid that may indicate the presence of *E. angustifolia*, and the presence of cichoric acid and caftaric acid in the other samples confirmed the presence of *E. purpurea*, however sample #49 do not allow an accurate estimation. Sample #48 labelled as including *E. purpurea* and *E. angustifolia* was also a glycerinated solution. Sample #50 labelled as including *E. pallida* confirms the presence of this species.

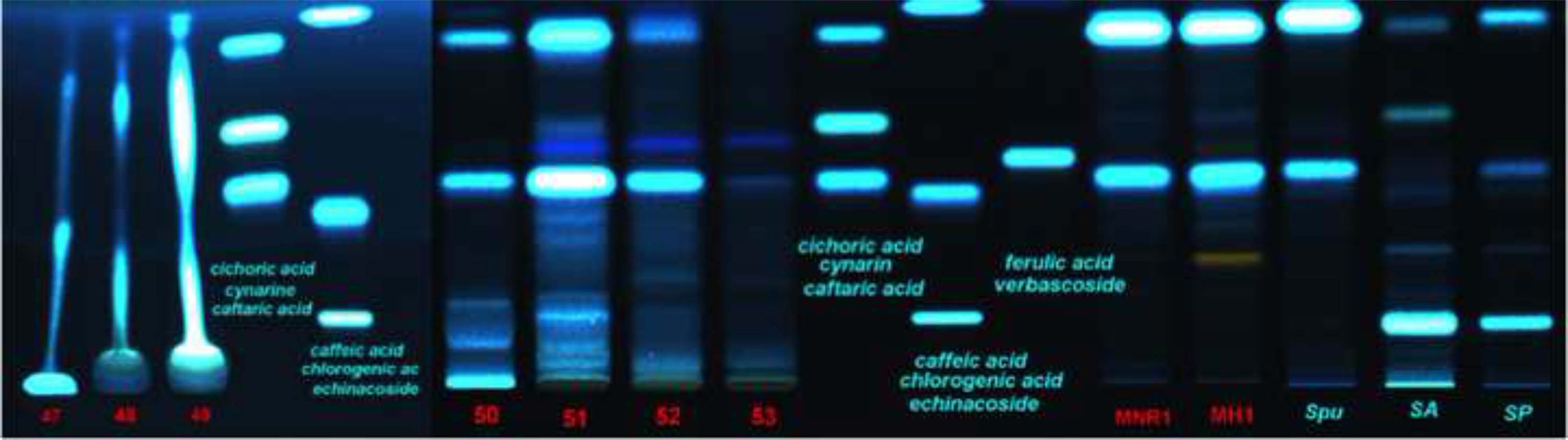

### 3.2 DNA metabarcoding

All 53 samples had detectable DNA concentrations but the results varied from sample to sample. Fifteen samples had a concentration lower than 0.1 ng/μl, 26 samples ranging from 0.1 to 10 ng/μl, and twelve higher than 10 ng/μl. No correlation between obtained DNA concentration and the substrate type of the extracted product was observed (Table S5).

The raw data before demultiplexing consisted of 12,190,865 sequences, with an average of 115,008 sequences per sample for each marker. After applying our trimming and filtering quality criteria, 38 herbal products (72 %) were retained, and they were used for further analysis (Table S6). Fifteen products, including three herbal teas (2, 7, 12) three capsules (19, 21, 29), five tablets (35, 36, 40, 43, 44) and four extracts (47, 50, 51, 53) did not yield reads or MOTUs after applying the quality filtering criteria and were excluded from further analyses. A total of 305,018 sequences passed the trimming and filtering quality criteria (2.5 *%* of reads), including 79,918 nrITS1 reads and 225,100 nrITS2 reads (Table S6). The MOTUs were formed using a 99 % similarity clustering threshold, and 2,529 MOTUs that contained minimum 10 reads were retained and further identified using BLAST as 83 different species (Table S7). For nrITS1 we detected a total of 60 different species and 37 species on only nrITS2 (Table S7). For both, nrITS1 and nrITS2, the number of species detected per sample ranged from 1 to 19, with an average of 5 species per sample.

**Figure 2.**
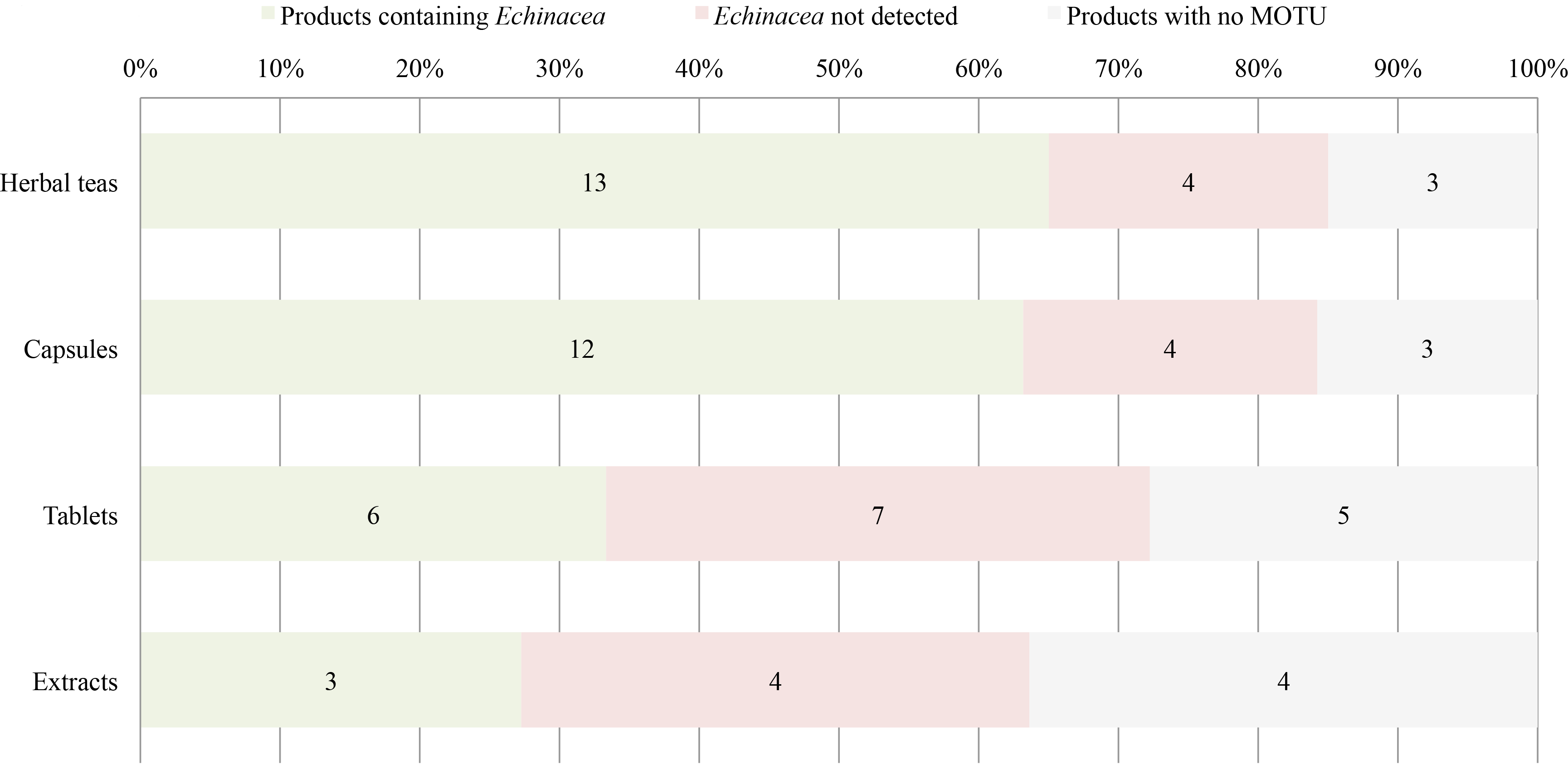
Detection of *Echinacea* sp. using DNA metabarcoding within the herbal products per category of pharmaceutical form.

The targeted *Echinacea* species were detected in 34 out of 38 products (89 %) of the retained samples (Figure 2 and Table S8). Twenty-four (89 %) out of 27 analysed products labelled as including *Echinacea* sp. as the single ingredient, contained at least one *Echinacea* species. However only seven (26 %) contained exclusively *Echinacea* species, whereas the other 20 contained several other species, likely contaminants. Ten (91 %) out of 11 analysed products labelled as including a mixture of *Echinacea* sp. and other plant species, contained *Echinacea* sp. but all of them showed considerable discrepancies between ingredients listed on the label and the ones detected. The overall ingredient fidelity, meaning the proportion of the number of species indicated on the product label and the entire species diversity, as detected by DNA metabarcoding, was 89 % for single ingredient products and 31 % for the multiple ingredient products. The overall ingredient fidelity for all products was 43 %.

Per formulation category the DNA metabarcoding yielded the following results (cf. Figure 2 and Table S8): In thirteen out of 17 samples the presence of *Echinacea* sp. was confirmed, and in the remaining four samples, three did not yield MOTUs that passed the quality filtering criteria. The capsules contained *Echinacea* sp. in twelve out of 16 samples, and identification was not possible in four of the products because these did not yield MOTUs that passed the quality filtering criteria. The herbal tables contained *Echinacea* sp. in only six out of 13 samples, and in the remaining seven samples five did not yield MOTUs that passed the quality filtering criteria. The extracts contained *Echinacea* sp. in only three out of seven samples, and the remaining four did not yield MOTUs that passed the quality filtering criteria.

## 4. Discussion

The quality and authenticity of herbal products have direct impacts on their safety. Quality control must include a series of procedures to ensure the identity of the used raw materials and screening of target compounds along the value chain. Authentication assays in herbal production must discriminate potential adulterants and/or substitutes. However, despite the existence of well-established and widely accepted analytical methods recommended in the regulatory guidelines (EDQM, 2014; EMA, 2006) for herbal product quality assessment, their resolution and efficacy can be impeded by various factors. First, the finished herbal products are often highly processed with numerous ingredients, and even if these analytical methods are accurate in detecting specific target compounds, they have limited efficiency in detecting infrageneric substitution and do not yield any information on other plant ingredients in the products (Rossi Forim et al., 2015). Morphology based taxonomic methods are equally impeded by highly processed herbal products as these often constitute of finely powdered materials (Zhao et al., 2006).

In this study, we combined HPTLC and DNA metabarcoding. HPTLC is an more automated and reproducible form of thin-layer chromatography (TLC), and with a better separation and detection of the compounds that can be successfully used in quality control of raw materials and finished herbal products (Reich and Widmer, 2008). Reich et al. (2008) summarized a procedure for HPTLC based identification of *Echinacea* and other species, which was applied on 53 herbal products in this study, including herbal tea, capsules, tablets and tinctures. The results showed that each of the three targeted *Echinacea* species have distinguishable chromatograms within most of the herbal products (Figure 1). The main limitation of the HPLTC assay is the inability to offer insights on the presence of other plant species in the herbal product. HPTLC is less suitable to for the analysis of volatile and certain sensitive samples (Morlock and Schwack, 2010). Combination of HPTLC with methods, such us mass spectrometry (MS), ultraviolet-visible (UV-VIS) and infrared spectrometry (IR) or Fourier-transformed infrared spectroscopy (FTIR) offer further possibilities for analytical refinement in the analysis of herbal products by increasing the amount of qualitative and quantitative information. In summary, HPTLC is not the most adequate method for detection of substitution and adulteration within marketed herbal products, but it is a powerful and cost-effective method to identify specific chemotaxonomic markers, and thus applicable in the quality control of the derived herbal products.

DNA barcoding and metabarcoding are not yet validated for use in a regulatory context of quality control (Agapouda et al., 2017), but there are several studies advocating its usefulness for herbal product authentication and pharmacovigilance (Cheng et al., 2014; Coghlan et al., 2012; Ivanova et al., 2016; Newmaster et al., 2013; Raclariu et al., 2017b, 2017a). The DNA metabarcoding results in this study show that the presence of *Echinacea* sp. was detected in 34 (89 %) out of 38 sequenced and retained samples. However, the results showed a lack of discriminatory resolution at the species level likely due to the low level of molecular divergence (Flagel et al., 2008; Zhang et al., 2017). Furthermore, most of the retained samples revealed a high level of discrepancy with most products not containing all the species listed on the label, but rather other off-label species. For all products, only 43 % of the species indicated on the product’s label were detected species using DNA metabarcoding. Here we need to mention that the general monograph number 1433 on ‘Herbal drugs’ of the European Pharmacopoeia allows up to 2 *%* foreign matter (EDQM, 2014), and that DNA metabarcoding is not a quantitative method that can be used to check if the contaminants are within this allowed range. As suggested by previous studies (Ivanova et al., 2016; Raclariu et al., 2017a), the results related to the authentication of herbal products using DNA metabarcoding need to focus primarily on checking the presence of the labeled ingredients and contaminants. The presence of non-listed species may be explained by various factors, including but not limited to the deliberate adulteration and unintentional substitution, that may occur starting from the early stage of the supply chain (i.e., cultivation, transport, storage), to the manufacturing process and the commercialization of the final products. DNA metabarcoding is a highly sensitive method and even traces of DNA, from grains of pollen from anemophilous species, for instance, or from plant dust in the entire manufacturing process that may accidentally contaminate the product, can be detected and identified (Raclariu et al., 2017b, 2017a). Several factors may influence the accuracy of the final results, starting with the first steps in processing the raw material to the final data analysis and interpretation. Extraction procedures, barcoding markers, primers, PCR amplification bias, high throughput sequencing library preparation, sequencing platform and trimming and filtering quality thresholds, clustering, and molecular identification algorithm all influence the final results (Pawluczyk et al., 2015; Staats et al., 2016). Moreover, public sequence reference databases, such as NCBI Genbank, pose significant challenges for reliable taxonomic affiliations, due to incorrectly identified or missing reference sequences and high levels of missing data (Hinchliff and Smith, 2014). However, at least one reference sequence of almost each genus of the known land plant is represented in NCBI GenBank (Hinchliff and Smith, 2014) and thus the use of DNA metabarcoding should be possible at least for identification at higher taxonomic levels.

As molecular genetics and sequencing technology advances, the use of DNA metabarcoding is promising for large-scale authentication of herbal products and other mixtures of economical importance. Standardization is required before DNA metabarcoding can be implemented as a routine analytical approach (Agapouda et al., 2017; Staats et al., 2016). Although, DNA metabarcoding is beneficial to authenticate species and study species diversity in complex mixture, it does not provide other essential information on target compound presence and concentration or the presence of chemical contaminations such as heavy metals, allergenic dyes and synthetic pharmaceuticals.

## 5. Conclusions

The increasing use of herbal medicines needs to be accompanied by an enhanced method for comprehensive quality control that are adequate for all the stages of the supply chain, starting from the cultivation of the raw material to the marketed herbal product. The results of our study show that HPTLC is a reliable analytical tool for routine use to identify and distinguish *Echinacea* species in herbal products. It allows for better separation and a course quantification of chemical constituents, but it has a limited resolution in detecting the presence of other species within the product. Here, we show that this limitation can be overcome through the complementary use of DNA metabarcoding that simultaneously confirms both the presence of the target species and of all other species, in even highly processed and multi-ingredient herbal products. Advances in sequencing technology advances make the use of DNA metabarcoding promising for large-scale authentication of herbal products. Nevertheless, standardization is required before it can be implemented as a routine complementary analytical method for regulatory quality control and herbal pharmacovigilance.

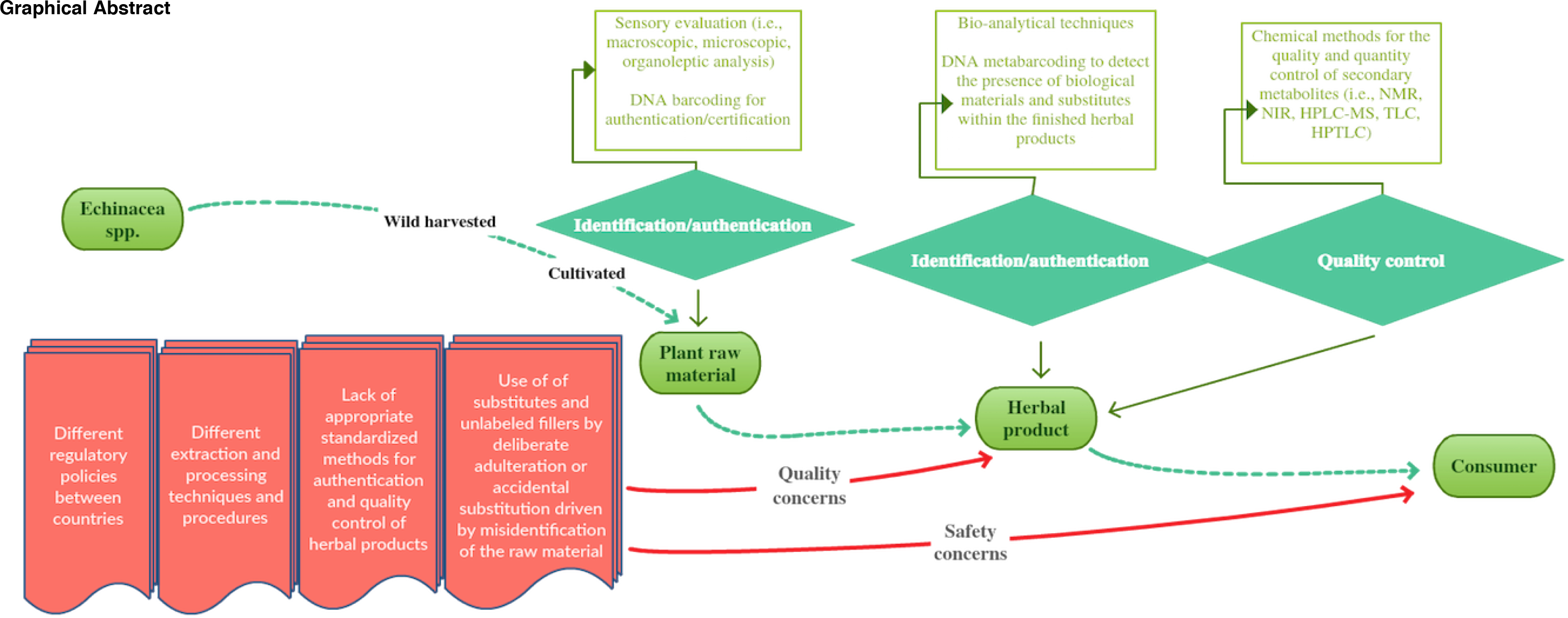

## Acknowledgments

The authors wish to thank the NHM head engineers Birgitte Lisbeth Graae Thorbek and Audun Schrøder-Nielsen and the DNA lab manager Jarl Andreas Anmarkrud for the technical help and assistance with the genetic analyses. Iulia Chilat and Elena Iacob are acknowledged for their technical help and assistance with the phytochemical analyses.

## Supporting information

Ion-Torrent amplicon read data is deposited in DRYAD [doi pending acceptance]

## Appendix. Supplementary materials

**Table S1** Use of medicinal *Echinacea* spp.

**Table S2** The main chemical constituents of **Echinacea** spp.

**Table S3** Information about the herbal products.

**Table S4** HPTLC results.

**Table S5** DNA and amplicon concentrations.

**Table S6** Overview of the results for 99 % clustering thresholds.

**Table S7** HTS reads and identified MOTUs per product.

**Table S8** Fidelity rate of the herbal products.

